# RNAseq analysis reveals the recurrent loss of heterozygosity in lung cancer and associated transcription patterns

**DOI:** 10.64898/2026.01.19.698133

**Authors:** Ruslan Gumerov, Wangzhen He, Phong Luong, Filippo Dall’Olio, Yegor Vassetzky, Anna Schwager

## Abstract

A key limitation in cancer transcriptomics is the lack of accompanying genomic profiling such as whole-genome sequencing (WGS) or copy number alteration (CNA) data. Here we address this by showing that RNA-seq alone can be used to infer chromosomal aberrations and identify biologically meaningful patterns in lung cancer. Through a large-scale meta-analysis of publicly available RNA-seq datasets from non-small cell lung cancer (NSCLC), small cell lung cancer (SCLC), and matched controls, we reconstructed large scale CNA profiles and identified deletions in 3p, 9p, and 17p as the most frequent genomic events. Validation against paired WGS data confirmed a high degree of accuracy for RNA-seq-based inference. Our analyses revealed that while deletion-associated transcriptional heterogeneity exists, approximately 25% of differentially expressed genes were shared across all three deletion classes, indicating a conserved oncogenic program in lung cancer. Enrichment analysis linked these shared genes to pathways governing cell division, DNA replication, and extracellular matrix organization, while deletion-specific effects reflected disruption of tumor suppressor pathways, notably p53 signaling in 17p-deleted tumors, leading to deregulation of NOS2 and PLOD2. Integrating gene-level expression data, we identified both shared and deletion-specific biomarkers: *TPX2* was consistently overexpressed across all deletion groups, while *PTPRZ1* and *CLDN9* were uniquely associated with del3p and del9p, respectively. Experimental validation in lung cancer cell lines confirmed these predictions, particularly the upregulation of *CLDN9* in del9p carriers. By demonstrating that RNA-seq data can capture large-scale chromosomal events and reveal their transcriptional consequences, this study establishes an efficient framework for genomic inference and biomarker discovery, introducing *CLDN9* as a novel, deletion-specific marker with potential prognostic and therapeutic value for 9p-deleted lung cancers.

---

RNAseq generates exabytes of data per year; many of these sequencing results concern cancer cells. In most cases, RNAseq analysis is not accompanied by WGS or CNA analysis, rendering the data incomplete and difficult to interpret in terms of precision cancer medicine. These data can potentially provide information beyond gene activity, including, e.g. annotation of SNPs based on sequencing data or short indel discovery, although the quality of indel annotation is quite low, particularly for large-scale CNAs that drive cancer progression. A number of algorithms for inference of large-scale CNAs from RNA-seq data were developed, including those based on beta-allelic frequency profiling and smoothing [1], though applications of these methodologies to real-world data still remain limited. In the current work, we present the results of an independent large-scale meta-analysis of publicly available bulk RNA-seq datasets of NSCLC, SCLC, and matching control samples. Characterization of the arm-level inferred deletion profiles of individual RNA-seq samples and benchmarking against reference WGS-based profiles reveals relative consistency of the inferred profiles with the reference data. We then focused on the analysis of specific CNAs subgroups (3p, 9p and 17p deletions) and revealed the associated transcriptional heterogeneity. Finally, we generated a list of novel deletion-specific biomarkers and provided validation *in vitro* using cancer cell lines carrying the deletions of interest, demonstrating the potential of *CLDN9* as a biomarker for del9p-carrying lung cancer subtypes.

## Indirect CNA inference and deletion annotation

We conducted a comprehensive analysis of publicly available lung cancer RNA-seq datasets and included in our study only datasets containing both control and cancer samples, with control samples consisting of adjacent normal tissues, although the number of controls did not always match the number of cancer samples. Our final analysis incorporated several publicly available RNA-seq lung cancer datasets (**Figure 1B**) [2–7]. All samples were reprocessed from raw .fastq files using the nf-core/rnaseq pipeline (see **Supplementary Methods**). TPM counts derived from tumor samples were subsequently utilized for CNA inference using CaSpER software (see **Figure 1A; Supplementary Methods**). All cancer samples included in the analysis were clinically classified as NSCLC, with the exception of the dataset SRP045225, which originated from SCLC patients. Following the deletion inference, the samples were categorized based on the presence or absence of specific CNAs (**Table S3**). We’ve identified del3p, del9p and del17p were the most common CNAs in NSCLC and SCLC patients (**Figure S1A**), in agreement with published data [8], suggesting the potential early-stage driving genetic events. Assuming that mentioned deletions represent driver aberration subtypes, we restricted our set with only samples carrying mentioned aberrations, resulting in an overall pool of 317 tumor samples.

**Figure 1.**
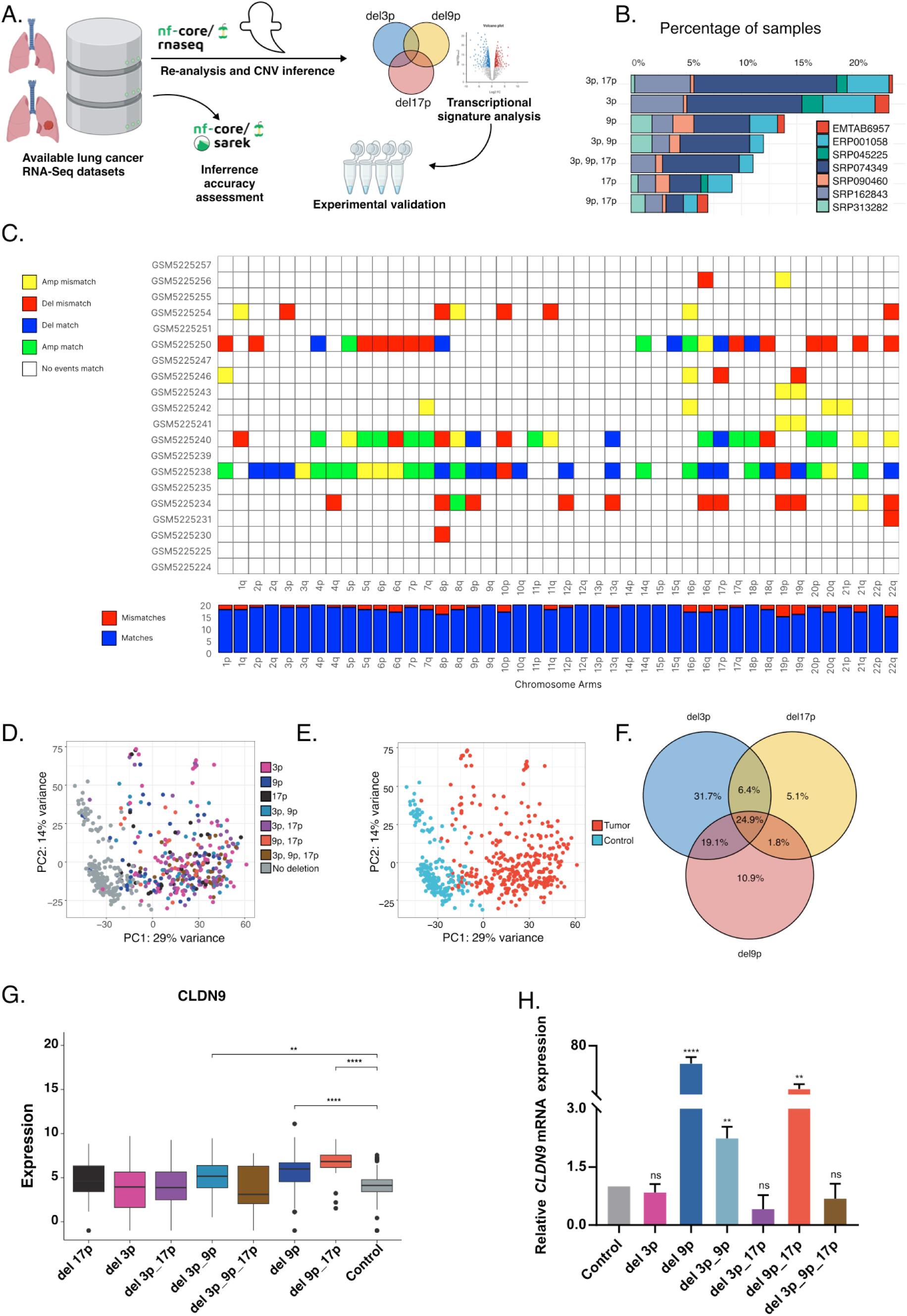
RNA-seq–based inference of chromosome arm-level events and associated transcriptional changes in lung cancer. **(A)** Schematic overview of the analysis workflow. Public lung cancer RNA-seq datasets were re-analyzed for gene expression quantification and chromosome arm–level copy number variation inference. Inferred CNV profiles were benchmarked against matched whole-genome sequencing (WGS) data where available, followed by transcriptional signature analysis and experimental validation. **(B)** Percentage of samples carrying chromosome arm-level deletions (3p, 9p, 17p, and their combinations) in analyzed RNA-seq datasets. Percentages are calculated relative to the total number of samples included in the analysis. Detailed information on inference methodology is provided in supplementary methods. **(C)** Comparison of RNA-seq–inferred CNVs vs. WGS-derived CNVs. Each row represents a sample and each column a chromosome arm. Colors indicate match or mismatch inference events for deletion or amplification calls. The bar plot below summarizes the number of mismatch and match call per chromosome arm. **(D, E)** Principal component analysis (PCA) of log-normalized gene expression profiles, colored by inferred deletion class **(D)** or sample type **(E)**. The proportion of explained variance shown on each axis. **(F)** Venn diagram showing overlap of differentially expressed genes (DEGs) associated with deletions of chromosome arms 3p, 9p, and 17p. DEGs were identified using Wilcoxon rank-sum tests comparing each deletion group to control samples, with Benjamini–Hochberg (BH) correction (adjusted *p* < 0.05). **(G)** *CLDN9* expression (log-normalized RNA-seq counts) across deletion groups and controls. Boxplots show median and interquartile range. Statistical significance was assessed using pairwise Wilcoxon rank-sum tests with BH correction (*p* < 0.05 *, < 0.01 **, < 0.001 ***, < 0.0001 ****). **(H)** Relative *CLDN9* mRNA expression measured by quantitative RT-PCR, normalized to a reference gene (*GAPDH*) and shown relative to controls. Statistical significance was assessed using One-way ANOVA test (ns, non-significant, p < 0.001 ***, < 0.0001 ****).

## Validation with the WGS data

The initial annotation of chromosomal events was inferred from RNA-Seq data utilizing the CaSpER software [9]. We next performed a validation study to assess the accuracy of deletion inference using the datasets with paired WGS samples from control and tumor samples, matched with the corresponding RNA-Seq data. The data was processed using the nf-core/sarek pipeline [10], with CNVKit option for copy number inference using the same thresholds for reference large-scale events annotation as above (see **Supplementary Methods**). For amplification and deletion combined, we achieved TPR of 68% and FPR of 2.1%, while maintaining the unbiased distribution of false calls. For amplification events, the results of CNA inference were slightly better than for deletions (TPR 75% and FPR 2.1% vs. TPR 61% and FPR 3%, respectively) (**Figure 1C, Table S3**). Altogether, in our study the inference methodology achieved the accuracy comparable to that of the original paper [10].

## Analysis of transcriptional signatures reveals deletion-associated heterogeneity

Lung cancer is associated with CNAs, but due to lack of paired RNAseq/WGS data, it has so far been difficult to analyze deletion signatures in NSCLC and SCLC. These transcriptional signatures of CNAs can reveal the etiology of lung cancer and provide biomarkers or treatment options.PCA analysis of transcriptional signatures revealed a separation between tumor and control samples groups along the PC1 (**Figure 1E**). The dispersion along the second principal component may be partially attributed to histological heterogeneity, caused by the distinction between the samples from the SCLC and NSCLC datasets (**Figure S1B**). PCA analysis of deletion-specific expression signatures for first 3 components (**Figures 1B, S1C**) did not reveal differences suggesting a degree of transcriptional similarity across different deletion profiles.

We next analyzed differentially expressed genes in samples carrying selected deletions (del3p, del9p, del17p) vs. the control group. Interestingly, we observed a largely conservative and co-regulated expression profile across all three deletion groups among differentially expressed genes: in all sets, a core of 24.9% genes was common to all three CNAs (**Figure 1F**). GO enrichment analysis performed on the common subset of genes revealed an enrichment in classical cancer-related pathways. Upregulated gene groups were associated with cell division, DNA replication, and extracellular matrix (ECM) organization, while downregulated gene groups were enriched for MAPK cascade, wound healing, and anatomical structure regulation pathways (**Table S3**). A comprehensive list of enriched pathways and their corresponding network plots are provided in (**Figure S2A, Table S3**). This observation can be attributed to the functional role of tumor suppressors as common cancer-associated deletions typically harbor genes that prevent malignant phenotype development through oncogene suppression. Removal of oncosuppressors leads to the development of a consistent cancer phenotype with conservative transcriptional signatures.

Importantly, a number of pathways that exhibited significant upregulation in deletion-specific groups demonstrated mechanistic connections to the deletions. For example, the amino-acid metabolic process term was selectively enriched in 17p deletion carriers (**Table S3, Figure S2B**) that are predominantly characterized by p53 loss and disruption of the p53 signaling cascade, with *NOS2* representing one of the downstream transcriptional targets regulated by p53. A direct link between p53 and transcriptional suppression of NOS2 exists [11] and NOS2 overexpression promotes oncogenic phenotypes [12]. *PLOD2* is another gene within the amino acid metabolism pathway that is specifically upregulated in cases with 17p deletion. Its expression is linked to disruption of the p53 pathway, as loss of p53 promotes HIF-1α-mediated upregulation of *PLOD2*, contributing to malignant phenotypes[13,14]. Despite the inherent constraints of indirect deletion inference methods, this analysis demonstrates their critical utility in uncovering biologically significant mechanisms underlying cancer phenotype development, even at a fine-grained, single-deletion level of analysis.

## Identification of deletion-agnostic and deletion-specific markers

We next performed gene-level differential expression analysis to identify genes differentially enriched across all of the deletion-carrier groups, as well as within each of the deletion groups separately, aiming to identify potential novel prognostic biomarkers. We have identified a group of candidate biomarkers with low to moderate prior literature support, representing promising targets for subsequent in vitro experimental validation.

One example is *TPX2,* which was significantly overexpressed in all of the deletion groups (**Figure S2C)**. TPX2 plays a critical role in microtubule dynamics regulation and cell division [15]. Overexpression of *TPX2* is a well-known trait of various cancers, including NSCLC [16,17].

We identified PTPRZ1 (**Figure S2C)** as a potential del3p-specific marker. In healthy tissues, *PTPRZ1* gene is expressed only in the neural system, while in various types of cancer, *PTPRZ1* expression may be either up- or down-regulated. In human gliomas, *PTPRZ1* was associated with poor prognosis, presumably via *PTN-PTPRZ1* axis [18]. *PTPRZ1* knock-down reduces proliferation in the glioma xenograft model [19] while overexpression of *PTPRZ1* in breast cancer is associated with chemotherapy resistance. Chemotherapy-driven increases in the CDKN1A/PTN/PTPRZ1 axis promote chemoresistance by activating the NF-κB pathway in breast cancer cells [20]. In lung cancer, *PTPRZ1* overexpression is detected by IHC inSCLC samples [21], but not in primary lung cancer tissues and brain metastases [22]*. PTPRZ1* knockout mice exhibit increased angiogenesis and elevated cell proliferation in induced lung adenocarcinoma models, presumably via *PTPRZ1-cMet* axis [23].

*DSC3* gene was also differentially expressed in del3p carriers except del3p, del9p (**Figure S2C**). The *DSC3* gene product, desmocollin 3, is a tumor suppressor gene and p53 target [24]. DSC3 is also overexpressed in lung squamous cell carcinoma relative to adenocarcinoma [25,26].

*HAPLN1* gene was specifically upregulated across all 17p-deletion-carrier groups, except the 9p, 17p. (**Figure S2C**). The protein product of *HAPLN1*, Hyaluronan And Proteoglycan Link Protein 1, maintains the extracellular matrix function by stabilizing aggregates of proteoglycan monomers with hyaluronic acid [30,31]. *HAPLN1* is associated with progression and metastasis in different cancer types, including lung cancer [32].

Finally, we proposed *CLDN9* as a del9p-specific marker; it was overexpressed in all 9p deletion carrier groups, with the exception of concurrent 3p, 9p, and 17p deletions (**Figure 1G**). *CLDN9* affects the survival levels and chemoresistance in breast cancer [27]. In gastric adenocarcinoma, overexpression of *CLDN9* enhances cell migration and proliferation [28]. *CLDN9* specifically, was associated with cell motility and invasiveness in Lewis lung carcinoma cell model [29].

The proposed markers were experimentally validated using a series of lung cancer cell lines carrying specific CNA **(Table S1**) and a primary lung fibroblast cell line used as a control. Our RT-qPCR result showed that *TPX2* was only overexpressed in del 9p and del 3p_17p groups (**Figure S2D**); *PTPRZ1* was significantly overexpressed in del 3p_17p and del 3p_9p_17p carriers (**Figure S2D**); *DSC3* was overexpressed in del 3p and del 3p_17p carrier groups and slightly underexpressed in del 3p_9p (**Figure S2D**); *CLDN9* was overexpressed in three all del_9p carrier groups except del 3p_9p_17p (**Figure 1H**).

In conclusion, our work establishes RNA-seq as a practical surrogate for genomic CNA profiling in lung cancer, revealing that transcriptomic data can uncover chromosomal aberration patterns and identify subtype-specific biomarkers. The methodology in this study was successfully applied with matched control samples and might be potentially adapted to reference-based approaches for larger datasets with unavailable controls. By linking inferred deletions to biologically coherent gene expression and validating novel markers like CLDN9, this study pioneers a powerful integrative framework for molecular stratification and biomarker discovery in precision oncology.

## List of Abbreviations

BH: Benjamini-Hochberg statistical method.
CNA: Copy number alteration.
CNV: Copy number variation.
CNVKit: Copy Number Variation Kit.
CLDN9: Claudin 9.
DNA: Deoxyribonucleic acid.
DSC3: Desmocollin 3.
ECM: Extracellular matrix.
FPR: False positive rate.
GO: Gene Ontology.
HAPLN1: Hyaluronan and Proteoglycan Link Protein 1.
HIF-1α: Hypoxia-inducible factor 1-alpha.
IHC: Immunohistochemistry.
Indel: Insertion or deletion.
LUAD: Lung adenocarcinoma.
MAPK: Mitogen-activated protein kinase.
NF-κB: Nuclear factor kappa-light-chain-enhancer of activated B cells.
NOS2: Nitric oxide synthase 2.
NSCLC: Non-small cell lung cancer.
PCA: Principal component analysis.
PLOD2: Procollagen-lysine, 2-oxoglutarate 5-dioxygenase 2.
PTN: Pleiotrophin.
PTPRZ1: Protein tyrosine phosphatase receptor type Z1.
RNA-seq: RNA sequencing.
RT-qPCR: Reverse transcription quantitative polymerase chain reaction.
SCLC: Small cell lung cancer.
SNP: Single nucleotide polymorphism.
TPM: Transcripts per million.
TPR: True positive rate.
TPX2: Targeting protein for Xenopus kinesin-like protein 2.
WGS: Whole genome sequencing.

## Acknowledgements

This work was funded by the Ligue Nationale Contre le Cancer, Gustave Roussy Emergence program and the IDB RAS Government basic research program (0088-2024-0010).

## Author contributions

Conceptualization, Y.V. and A.S.; Methodology, R.G., W.H. and A.S. Data analysis and curation, R.G., W.H., Y.V. and A.S.; Investigation, R.G., W.H., Y.V. and A.S.; Writing – Original draft: R.G. and Y.V.; Writing - Review & editing: R.G., W.H., F.DO., Y.V. and A.S.; Visualization, R.G., W.H. and A.S.; Funding acquisition, Y. V. and F.DO

## Supplementary data

### MATERIALS AND METHODS

#### Alignment and read quantification

Raw data from publicly available datasets GSE87340, GSE120622, GSE60052, GSE171415, GSE81089, E-MTAB-6957 and GSE40419 [2–7] were downloaded to the HPC cluster using sra-prefetch. For the read alignment and quantification task we used the netflow workflow management system (version 23.04.1) and the nf-core/rnaseq pipeline (version 3.12.0). We additionally specified TrimGalore for trimming adapters, STAR [33] for alignment and Salmon for quantifying the reads [34]. The obtained MultiQC report was investigated before the downstream analysis.

#### Reference WGS data annotation

Raw data from the phs001697 dataset [7] was used to perform the structural variant annotation for the samples from GSE171415 [7]. For the read alignment and quantification, we used the nextflow workflow management system (version 23.04.1) and the nf-core/sarek pipeline. For the annotation of the structural events, we used the CNVkit software [35], with the same thresholds that were used in the inference paper [9]. Thus, the event was annotated as deletion if the mean log ratio value was lower than −0.3, as an amplification if the mean log ratio value was higher than 0.3. We considered a deletion/amplification event to be arm-level if it exceeds one-third of a chromosomal arm’s length.

#### Structural variant inference

Variant inference and all downstream analyses were performed using R programming language. For the inference of structural variants we utilized CaSpER tool [9]. We performed BAF generation using the BAFExtract subprogram and then proceeded with CaSpER annotation with expression cutoff for lowly expressed genes equal to 3. Then we performed gene-based CNV summarization and performed CNA annotation (**Figure 1A**). For deletion annotations, we used the following methodology: we obtained lists of genes included in the deletion signature of each of the analyzed deletions. We used *RBM6, SEMA3F, SEMA3B, GNAT1, NAA80, HYAL1, HYAL2, HYAL3, NPRL2, CYB561D2, TMEM115, CACNA2D2, HEMK1, CISH* genes as 3p21 signature genes; *IFNB1, IFNA21, IFNA7, IFNA10, IFNA16, IFNA17, IFNA14, IFNA2, IFNA8, IFNE, MTAP, CDKN2A* for 9p21 signature; *AIPL1, SLC13A5, XAF1, EIF5A, FGF11, POLR2A, SOX15, SHBG, ATP1B2, TP53, ALOXE for the 17p signature*. We considered a sample to have the desired deletion only in case if all of the signature genes were depleted, or all but one of them. For the visualization of obtained variant data we used a modified version of plotLargeScaleEvent function available in the project github (https://github.com/braaist/IGR_lung_cancer).

#### Differential expression analysis

We used the DESeq2 package to analyze differential expression. Salmon read counts were imported using the “tximport” library. The integrated object was analyzed using a dataset of origin as a covariate for differential expression analysis. Final LFC values for comparisons between samples with different deletions and controls were shrunken using “normal” shrinkage estimator, and visualized using “EnhancedVolcano” library. As a cutoff for the significance of differential expression in samples, we used adj.p.value < 0.001 and absolute LFC value greater than 2.5. VST counts were used for PCA analysis and expressional boxplots.

#### Functional enrichment

We used R libraries “enrichplot”, “DOSE” and “clusterProfiler” for functional enrichment. Ensembl gene identifiers obtained on the alignment stage were converted to the ENTREZID and analyzed using gseGO and enrichGO functions with Benjamini-Hochberg p value adjustment and with pvalueCutoff equals to 0.05.

#### Technical equipment

All data obtained in this article was generated using HPC cluster with slurm scheduler (IGR server Flamingo).

#### Cell culture

We used a series of lung cancer cell lines with specific CNAs (**Table S1**). A549 and NCI-H1299 were grown in Dulbecco’s Modified Eagles Medium (DMEM) (Gibco, USA) supplemented with 1% v/v Penicillin-Streptomycin Solution (PST) (Gibco, USA) and 10% v/v Foetal Bovine Serum (FBS) (Gibco, USA). The others were cultured in RPMI 1640 (Gibco, USA) supplemented with 1% v/v PST and 10% v/v FBS.

**Table S1.**
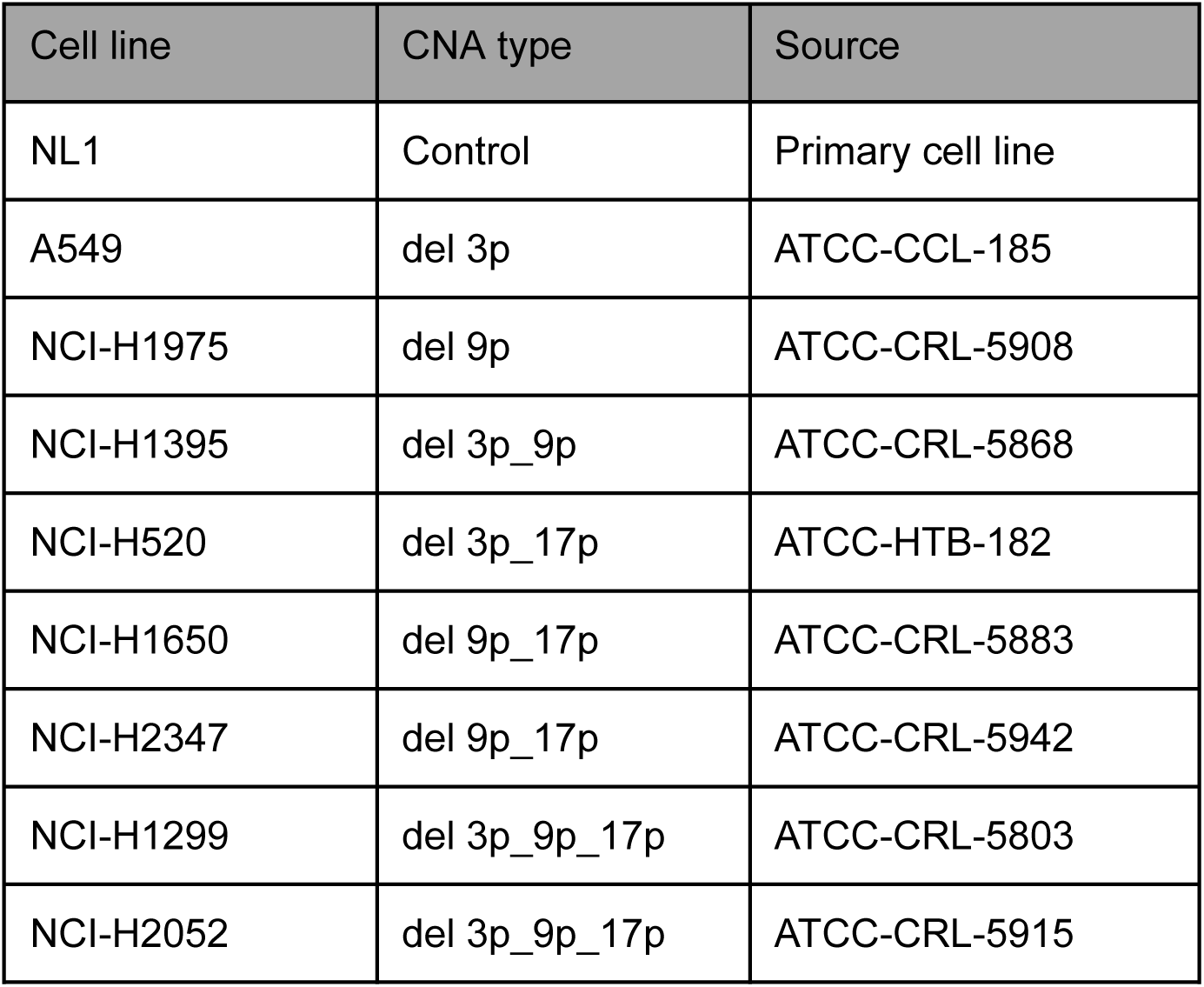
Lung cancer cell lines by CNA type.

| Cell line | CNA type | Source |
| --- | --- | --- |
| NL1 | Control | Primary cell line |
| A549 | del 3p | ATCC-CCL-185 |
| NCI-H1975 | del 9p | ATCC-CRL-5908 |
| NCI-H1395 | del 3p_9p | ATCC-CRL-5868 |
| NCI-H520 | del 3p_17p | ATCC-HTB-182 |
| NCI-H1650 | del 9p_17p | ATCC-CRL-5883 |
| NCI-H2347 | del 9p_17p | ATCC-CRL-5942 |
| NCI-H1299 | del 3p_9p_17p | ATCC-CRL-5803 |
| NCI-H2052 | del 3p_9p_17p | ATCC-CRL-5915 |

#### RT-qPCR

Total RNAs were extracted from adherent cells based on the protocol of Quick-RNA MiniPrep Kit (Zymo Research, USA) and RNA quantification was assessed using a NanoDrop spectrophotometer (Thermo Fisher, USA). Reverse transcription was carried out to obtain the correspondent cDNA using HiScript III RT SuperMix (Vazyme, China) and cDNAs were diluted five-fold prior to qPCR analysis. qPCR (10 μL reaction volume) was carried out using Taq Pro Universal SYBR (Vazyme, China) and Ct values were collected by FMR-5S Smart Quant System (Vazyme, China) with specific primers (**Table S2**).

**Table S2.**
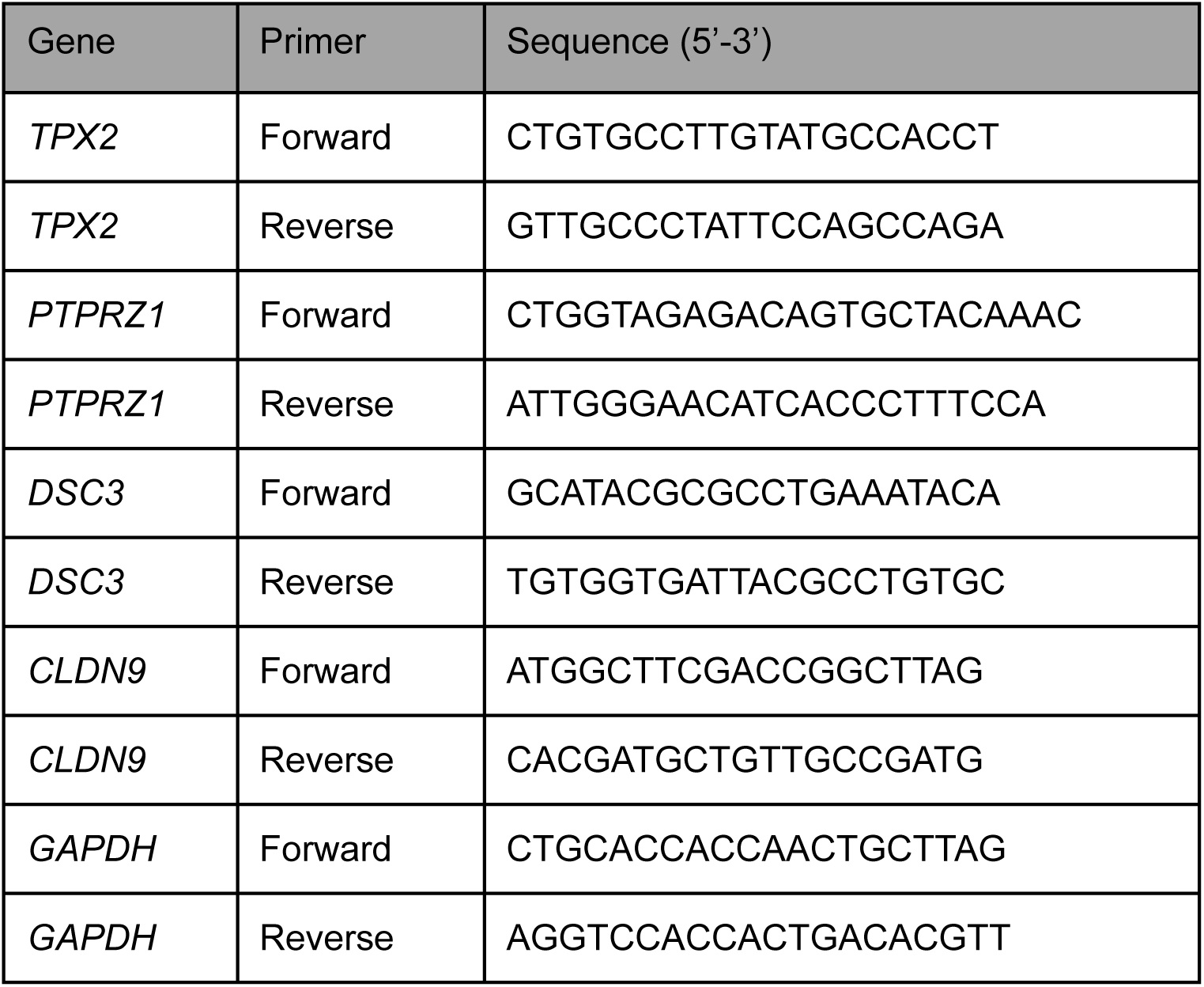
Primers used for qPCR.

## Data availability statement

All data and code generated during this project is available on the github project page (https://github.com/braaist/IGR_lung_cancer).

## Supplementary Figures

**Figure S1.**
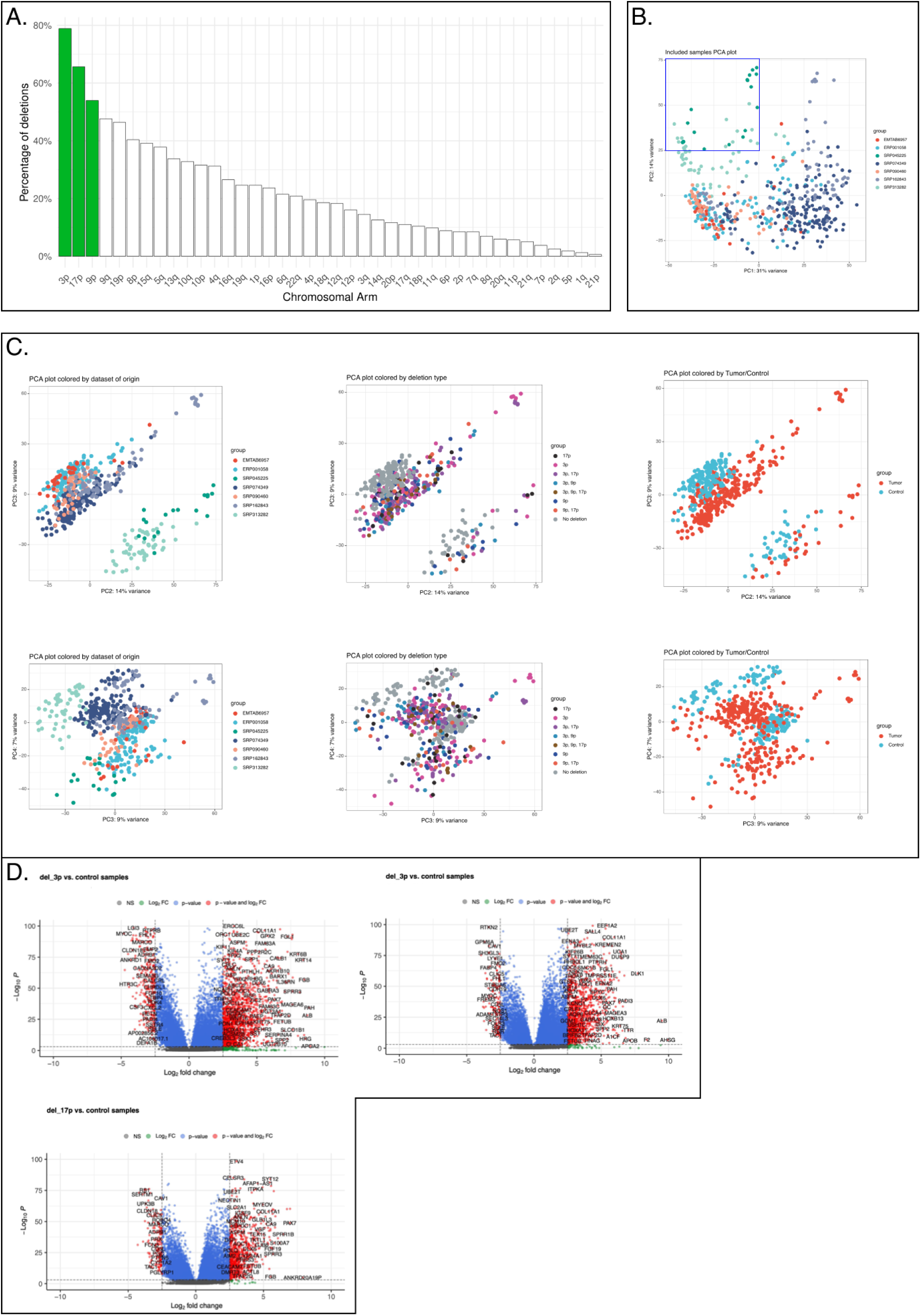
Chromosomal deletion landscape and transcriptional impact in lung cancer cohorts. **(A)** Distribution of inferred arm-level deletion profiles among tumor samples included in the analysis. Green bars highlight the three most frequent deletions (3p, 17p, 9p). Percentages represent the proportion of samples carrying each deletion relative to total analyzed samples. (**B**) PCA of gene expression profiles showing first and second principal components, colored by dataset of origin. The small cell lung cancer (SCLC) dataset SRP045225 is highlighted with a blue frame. Proportion of variance explained is shown on each axis. (**C**) PCA plots showing second, third and fourth principal components, colored by dataset of origin (left), deletion type (middle), and sample classification as tumor or control (right). Top row shows PC2 vs. PC3; bottom row shows PC2 vs. PC4. (**D**) Volcano plots of differential gene expression comparing samples with specific deletions (del_3p, del_3p, del_17p) versus control samples. X-axis shows log₂ fold change; Y-axis shows −log₁₀ p-value. Points are colored by significance: gray (not significant), green (log₂FC threshold only), blue (p-value threshold only), red (both p-value and log₂FC thresholds). Gene names are labeled for selected significantly differentially expressed genes. Log2 fold changes were shrunk using the “normal” shrinkage estimator from DESeq2. Genes were considered differentially expressed if they met dual thresholds: Benjamini-Hochberg adjusted p-value < 0.001 and absolute log2 fold change > 2.5.

**Figure S2:**
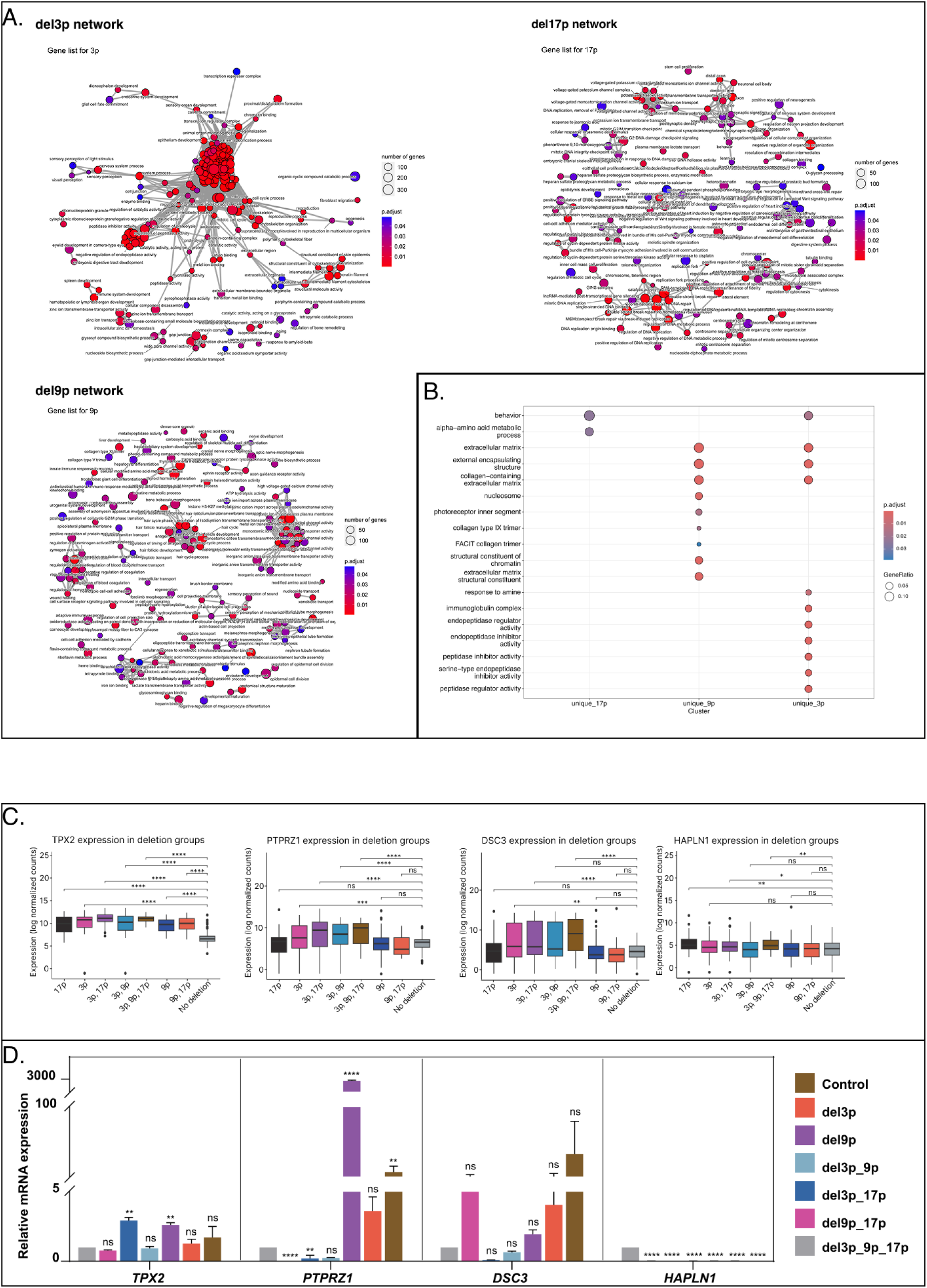
Functional enrichment analysis and expression validation of deletion-specific transcriptional signatures. **(A)** Network plot of Gene Ontology terms for genes uniquely differentially expressed in del_3p, del_9p, and del_17p samples compared to controls. Plots generated using clusterProfiler enrichGO with pairwise term similarity. Node size represents the number of genes associated with each GO term; node color indicates adjusted p-value. Only genes unique to each deletion type were included in the analysis. GO terms were filtered for minimum gene set size of 3 and maximum of 800, with Benjamini-Hochberg adjusted p-value < 0.05. (**B**) Dot plot showing deletion-specific GO term enrichment from compareCluster analysis. Dot size indicates gene ratio; dot color indicates adjusted p-value. Only terms with significant enrichment (adjusted p < 0.05) in at least one deletion group are shown. Only genes unique to each deletion type were included in the analysis. (**C**) Expression of candidate deletion-specific marker genes across deletion groups and controls. Box plots show log-normalized RNA-seq counts for TPX2, PTPRZ1, DSC3, and HAPLN1. Each box represents the interquartile range with median line; outliers shown as individual points. Statistical significance assessed using pairwise Wilcoxon rank-sum tests with Benjamini-Hochberg correction (ns = not significant, * p < 0.05, ** p < 0.01, *** p < 0.001, **** p < 0.0001). Colors correspond to deletion groups as defined in Figure 1. (**D**) Quantitative RT-PCR validation of candidate marker gene expression in distinct LOH-type lung cancer cell lines. Relative mRNA expression normalized to GAPDH reference gene and shown relative to NL1 control cell line. Bars represent mean ± SD from at least three biological replicates. Statistical significance assessed using One-way ANOVA testcomparing each deletion group to NL1 control (ns = not significant, ** p < 0.01,**** p < 0.0001).

